# Construction of a low-temperature degrading synthetic bacterial community of corn straw

**DOI:** 10.1101/2025.02.07.637170

**Authors:** Yi Fang, Jiaqi Li, Wanyi Zhang, xuhong Ye, Xiaolin Zhu Zhu, Yang Xie, Youzhi Feng, Yongjie Yu, Susu Yu, Li Zhang, Hongtao Zou

## Abstract

The stability of the cellulose structure in straw creates a challenge for short-term degradation in the natural environment, especially in the cold regions of north China. Therefore, to secure strains that can degrade straw at low temperatures, we selected soil samples from straw-returning land and used cellulose as the only carbon source. After conducting preliminary screening and further re-screening, we ultimately selected three strains that exhibited efficient degradation of corn straw at a temperature of 4 ℃. Subsequently, the synthetic bacterial community C was constructed using these strains: *Stenotrophomonas maltophilia*, *Flavobacterium johnsoniae* and *Pantoea rodasii*.

We conducted a 45-day solid-state and liquid-state fermentation experiment at low-temperature (12 ℃). The activity of lignocellulase was determined, and the enzyme production conditions were optimized. The results show that the liquid fermentation degradation rate of straw increased by 40 % compared with the control group in the natural state (without adding C). Similarly, the solid-state fermentation degradation rate of straw increased by 30 %, which was significantly faster than that of the control group. Finally, the enzyme production conditions of strain C were optimized as follows: culture time 96 h, inoculation amount 4 %, temperature 12 ℃, pH 6.0, and Alkaline Carboxymethyl Cellulase (CMCase) activity 24.34 U/mL. The microbial system C efficiently degrades straw at low temperature, which is valuable for improving the comprehensive utilization rate of straw and has a broad application prospect in improving soil fertility.

## 1. Introduction

As the main agricultural waste, straw returning is of great significance for improving soil carbon stocks, improving soil physical and chemical properties and impacting the soil ecological environment. (Chen et al., 2018) However, most of the corn straw is derived from plant cell walls, and the biopolymers formed by cellulose, hemicellulose and lignin are difficult to degrade, which brings great challenges to its natural decomposition in the field (J. Wang et al., 2019), especially in Northeast China, where the temperature is low in winter.

Traditional chemical treatment technology is difficult, and the associated pollution is large. The new biotechnology involves microorganisms with straw-degradation ability, which can make up for the shortcomings of traditional treatment technology, and prevent environmental pollution. (Chang et al., 2021) The microbial degradation process works through the production of various lignocellulosic hydrolases, including exonuclease, endoglucanase and β-glucosidase. Endoglucanase acts on the crystalline region of cellulose, which makes cellulose easy to hydrate. (Xue et al., 2017) Endoglucanase acts on the amorphous region of cellulose, which decomposes β-1,4 glycosidic bonds into cellobiose and other oligosaccharides, and then these decomposes into glucose by β-glucosidase (Wang Q. L. et al., 2004). Thus, microbes can decompose difficult-to-degrade lignocellulose into small molecules of glucose and improve the utilization efficiency of straw. The strains involved in the study are mainly actinomycetes, fungi and bacteria, of which fungi produce large amounts of cellulase enzymes, which play a dominant role in the straw degradation process. (Xu et al., 2011) However, some studies have pointed out that bacteria have stronger environmental adaptability and biochemical versatility than fungi (Lee et al., 2019).

Microbial degradation of straw is affected by many factors, among which temperature is the main one affecting the function of microbial degradation (A. J. Chang et al., 2012). In Northeast China, the natural degradation of a large amount of the agricultural by-product straw is a huge challenge. The undegraded straw not only affects the growth of crops in the second year, but also causes crop diseases and pests (Yang et al., 2020). Therefore, it is of great significance to seek microbial sources with cellulose-degradation ability at low temperature to accelerate straw decomposition and improve soil fertility in colder regions (Zheng et al., 2020). Composite microbial agents are more adaptable and better interact than single strains, and thus have a good degradation effect (Chen et al., 2019; Puentes Téllez & Salles, 2018). A variety of microorganisms interact with the degradation environment through their own enzymes to form a specific flora, and use their respective functions to achieve straw degradation (Ozbayram et al., 2018).The general method of constructing microbial inoculants is to use the traditional pure culture technology to screen a single high-efficiency strain, and then to construct a composite decomposing inoculant that produces high-efficiency cellulase through antagonism test, optimized combination technology and synergy between multiple microorganisms (Xu et al., 2011). For example, the composite microbial agent (CB) was constructed with *Phanerochaete* and *Chrysosporium*. After 20 days of degradation under the optimum conditions (32 ℃), the degradation rates of lignin, cellulose and hemicellulose of corn straw were 44 %, 31 % and 48 %, respectively, which enhanced the degradation rate of corn straw (Chu et al., 2021). Moreover, the complex microflora composed of *Clostridium*, *Acinetobacter*, *Bacteroides*, *Lysinibacillus* and *Dysgonomonas* had a high capacity to produce xylanase and carboxymethyl cellulase, and the enzyme activity was stable at pH 5.0-9.0. The feasibility of the complex microbial enzyme coordination mechanism for lignocellulose degradation was confirmed (Zhu et al., 2016).

Based on recent information, we finally screened three strains of bacteria with high cellulose-degradation ability at low temperature and constructed a synthetic bacterial community C. In this study, we further confirmed the feasibility of composite microbial enzyme coordination mechanism for lignocellulose degradation. By supplementing bacterial community C in the process of straw returning to the field, it provides microbial resources for accelerating the degradation of straw in winter and improving the soil fertility. At the same time, it also provides technical support for improving crop yield and preventing pests and diseases in the second year.

## 2. Materials and methods

### 2.1 Screening for corn straw degradation

The process of bacterial community construction comprised three main stages: sampling and culturing, primary screening, and re-screening. Soil and corn straw were collected from an experimental field with eight years of continuous corn planting in Saraki farmland, Shenyang city, China, where the samples were used the same day.

#### 2.1.1 Primary screening of corn straw degradation

Aliquots of 10 g sampling were added to sterilized distilled water (90 mL) in flasks. Flasks were shaken at 180 rpm for 30 min, and then left standing for 30 min (Shan et al., 2015). One mL of bacterial suspension was inoculated into 100 ml of Hutchison’s liquid medium (KH_2_PO_4_ 0.1 % (w/v), FeCl_3_ 0.001 % (w/v), MgSO_4_·7H_2_O 0.03 % (w/v), NaNO_3_ 0.25 % (w/v), CaCl_2_ 0.01 % (w/v), NaCl 0.01 % (w/v) and a filter paper strip was included), and placed at 20 °C (Zhang et al., 2002). The culture at the disintegration of the filter paper strip was picked up and transferred to the CMC liquid medium (CMC-Na 1.5 % (w/v), NH_4_NO_3_ 0.1 % (w/v), yeast extract 0.1 % (w/v), MgSO_4_ · 7H_2_O 0.05 % (w/v) and KH_2_PO_4_ 0.1 %(w/v)). The culture was shaken at 20 ° C and 180 rpm for three days. Then, the bacterial solution was gradient diluted from 10^-4^ to 10^-6^ under sterile conditions, 1 mL for each dilution, coated the CMC solid medium (2 % agar was added to the liquid medium). Each gradient set up three parallels and repeated purification and separation until a single strain was obtained.

#### 2.1.2 Re-screening of corn straw degradation

A single colony with obvious morphology on the plate was selected, and the activity of sodium carboxymethyl cellulose (CMCase) was determined by Congo red staining. The ratio of the diameter of the transparent circle (D) to the diameter of the colony (d) was recorded. In addition, we also determined the capacity of the strain to produce lignocellulose-degrading enzymes, including filter paper activity (FPA), endoglucanases (EG), and β-glucanases. In a continuous process, the degradation rate of corn straw by a microbial consortium under soil conditions was measured on the 45^th^-day. Finally, the purified cellulose-degrading monocultures were cultured on CMC solid medium for seven to 30 d at 20°C to 4°C in a gradient of 4°C, and the strains that could grow under the conditions of 4 to 20°C were selected.

### 2.2 Construction of a cellulose-degrading complex microbial system

First, we did antagonism tests between each strain. Finally, strains without an antagonistic reaction were mixed, numbered (A, B, C, D) and the composition of each combination was recorded. The strains were transferred to the new CMC liquid medium. Cultures were grown on an orbital shaker (180 r.p.m.) at 20 °C for 12 hours.

### 2.3 Identification of isolated bacterial strains

#### 2.3.1 Identification of colony morphology and physiological and biochemical characteristics

The selected single bacteria were inoculated on Luria Broth solid medium and cultured at 20 ° C for 48 h. The colony morphology of each strain was observed and a Gram staining was performed. The strains were identified by physiological and biochemical tests according to《Berger’s Manual of Bacterial Identification》and 《Manual of Systematic Identification of Common Bacteria》.

#### 2.3.2 Molecular biological identification of strains

In this experiment, the total DNA of a strain was extracted by the Tangen Bacterial Genome Extraction Kit (*Tangen Biochemical Technology, Beijing, China*), and amplified by primer sets 27F and 1492R for high-throughput sequencing. The PCR reaction was performed in 25 μL reaction mixture, including 2.5 μL 10 × buffer, 2 μL dNTPs (5 mM), 1.5 μL MgCl_2_ (1.5 mM), 1 μL forward primer (10 mM), 1 μL reverse primer (10 mM), 0.3 μL rTaq DNA polymerase (0.4 μL TaKaRa, Japan), 1 μL DNA template (approximately 50 ng of genomic community DNA) and 15.7 μL ddH2O. Then 30 cycles (95°C 5 min, 94°C 30s, 55°C 30, 72°C 1min), the final extension at 72 ° C for 10 min. At the end of the reaction, the samples with multiple bands were subjected to gel recovery of the target fragments using a gel recovery kit (Takara, Beijing, China). The PCR product was prepared using the pEASY-T1 Cloning Kit (Beijing, China), and the fragments of the above PCR product were inserted into the cloning vector to obtain the plasmid with the fragment to be tested. The amoA gene of total bacterial DNA of each strain was amplified by PCR and cloned by amoA-27F / amoA-1492R. The positive clones were identified by PCR, and the specific reference (Li et al., 2009). After the preparation, the positive clones were sent to Sangon Bioengineering (Shanghai, China) for sequencing. The sequencing primers were M13 Forward and M13 Reverse primers. One sequenced sequence of each strain was selected and compared in the GenBank database of NCBI, and the sequences of the model strains with high sequence similarity of homologous genes were downloaded.

### 2.4 Measurement of lignocellulolytic enzymes

The 5 mL bacterial suspension of the screened strain was cultured in liquid enzyme-producing medium for three days, and centrifuged (5000 r. p. m.) for 10 min. The supernatant was the crude enzyme solution. The DNS colorimetric method was slightly modified (Wang et al., 1998) to determine FPA, endoglucanase activity (CMCase) and β-glucosidase activity (β-Caes) of these strains and synthetic colonies. Measurement of OD was made at a wavelength of 540 nm, and the specific enzyme activity were calculated according to the standard curve. The specific enzyme activity reflects the rate of glucose production. The calculation was as follows:

C•10 / T•V

C is glucose content (μg), T is reaction time (min), and V is crude enzyme liquid volume (mL). Production of enzyme units (U) was defined by the ability to transform substrate into product each one per minute.

### 2.5 Determining degradation rate of corn straw lignocellulose

To verify the efficiency and practical application of the above selected straw-degrading bacteria, these bacteria were subjected to low-temperature solid-state fermentation and liquid-state fermentation at 12°C for 45 days, and the weight loss of straw was calculated.

#### 2.5.1 Liquid fermentation of straw-degrading bacteria

A total of 2 g dry corn straw and 5 mL bacterial suspension of low-temperature cellulose-degrading bacteria were added to 100 mL liquid fermentation nutrient solution (Peptone 0.2 % (w/v), MgSO_4_ · 7H_2_O 0.05 % (w/v), KH_2_PO_4_ 0.1 % (w/v), NaCl 0.05 % (w/v), CaCl_2_ 0.02 % (w/v) and yeast extract 0.05 % (w/v)). Each strain had three replicates and was cultured at 12° C and 120 rpm for 45 days. The precipitate was collected after centrifugation (6000 r. p. m.) of the culture solution for ten minutes, rinsed repeatedly by centrifugation with dilute acid, and finally rinsed by centrifugation with distilled water. Then it was dried to constant weight, to calculate the rate of weight loss of straw before and after the experiment.

#### 2.5.2 Solid-state fermentation of straw-degrading bacteria

Aliquots of 10 g dried corn straw and 2 mL bacterial suspension were added to 20 ml solid-state fermentation nutrient solution (the composition was the same as the liquid fermentation culture medium, and the concentration was increased five times), with three replicates for each strain. Culturing time was 45 d at 12°C and shaking every 5 days, to calculate the rate of weight loss of straw during the experiment.

### 2.6 Optimization of enzyme production in cellulosic complex strains

To evaluate the effects of different factors on the degradation of corn straw by different strains, the factor levels of each experimental group were designed as follows: strain inoculum (2%, 4%, 6%, 8%, 10%), temperature (4℃, 8℃, 12℃, 16℃, 20℃), training time (24 h, 48 h, 72 h, 96 h, 120 h) and pH (5.0, 6.0, 7.0, 8.0, 9.0). Each test group was cultivated in liquid fermentation medium (180 r. p. m.) for three days, and the CMC enzyme activity and OD540 were measured using the one without bacterial solution as a control (Gou et al., 2017).

## 3. Results

### 3.1. Selection of low-temperature cellulose-degrading strains

After the culture had been transferred for 20-30 generations, the filter paper strip steadily began to disintegrate, and some colonies grew in the contact part of the filter paper strips and the culture medium. The cellulose-degrading strains were isolated by the plate dilution method on CMC solid medium. According to the color, size and shape of the colony, 43 strains of single bacteria were screened and purified. Combined with Congo red identification test, 10 single strains were selected and numbered. Then, a temperature gradient of 20°C, 16°C, 12°C, 8°C and 4°C was set for the low-temperature induction of the strain. With the decrease of culture temperature, the growth rate of some test strains showed a slow trend, and some strains were unable to acclimate to the low temperature. Only strains X24, X26 and X37 continued to grow at 4 ° C to 20 ° C (Table S1). Finally, three strains (X24, X26, X37) with the function of degrading corn straw at low temperatures were selected.

### 3.2 Taxonomic composition of the selected bacterial

The sequence lengths of strains X24, X26 and X37 were 1450 bp, 1417 bp and 1443 bp, respectively. The results of NCBI sequence alignment showed that the 16 S rDNA sequence strains with the highest similarity to X24 belonged to *Stenotrophomonas*, and the homologous sequence similarity reached 99 % (Table S2). The sequence alignment results of strain X26 showed that the similarity between strain X26 and *Flavobacterium* was 99 % (Table S3). The sequence alignment of strain X37 showed that the similarity between strain X37 and *Pantoea* was 96 % (Table S4). Combined with the morphological observation and physiological and biochemical characteristics of the strains (Table 1), the strain X24 was preliminarily identified as *Stenotrophomonas maltophilia*, strain X26 was identified as *Flavobacterium johnsoniae* and strain X37 was identified as *Pantoea rodasii*.

**Table 1.** Physiological and biochemical test results of three strains.

| Pilot Project | X24 | X26 | X37 |
| --- | --- | --- | --- |
| Gram stain | - | - | - |
| VP test | + | + | - |
| OF test | + | + | + |
| Indole test | - | - | - |
| Gelatin liquefaction test | + | - | + |
| Catalase test | + | - | + |
| Oxidase test | + | + | + |
| Nitrate reduction test | - | + | + |
| Utilization of citrate | + | - | - |
| Methyl red test | + | + | + |
| Starch hydrolysis test | - | + | + |
| Motility | + | - | + |
| Lipase test | - | + | + |

Note: "+" means positive reaction and "-" means negative reaction.

### 3.3 Construction of a low temperature straw-degrading microbial community

Antagonism tests were carried out on strains X24, X26 and X37. Results show that the three strains did not antagonize each other, so the formation of a bacterial community could be carried out. Four bacterial communities were constructed and named A (X24 and X26), B (X24 and X37), C (X24, X26 and X37) and D (X26 and X37).

### 3.4 Lignocellulose-degrading enzymes activity

CMCase enzyme activity, FPA enzyme activity and β-glucosidase enzyme activity were determined for single strains X24, X26, X37 and composite strains A, B, C and D, respectively. The results show that the three single strains all contained enzymes that decomposed lignocellulose (Fig. 1). Among them, *Pantoea rodasii* had the highest enzyme activity, CMC enzyme activity was 21 U/ mL, FPA enzyme activity was 14 U / mL, and β-glucosidase enzyme activity was 12.3 U / mL. In the bacterial community, the enzyme activity of community C was significantly higher than that of other communities. The CMCase enzyme activity was 25 U / mL, the FPA enzyme activity was 18 U / mL, and the β-glucosidase enzyme activity was 15 U / mL. Indicating that the community C showed higher cellulose degradation ability than other test groups. Compared with a single strain, the four communities had higher cellulase activity.

**Fig. 1.**
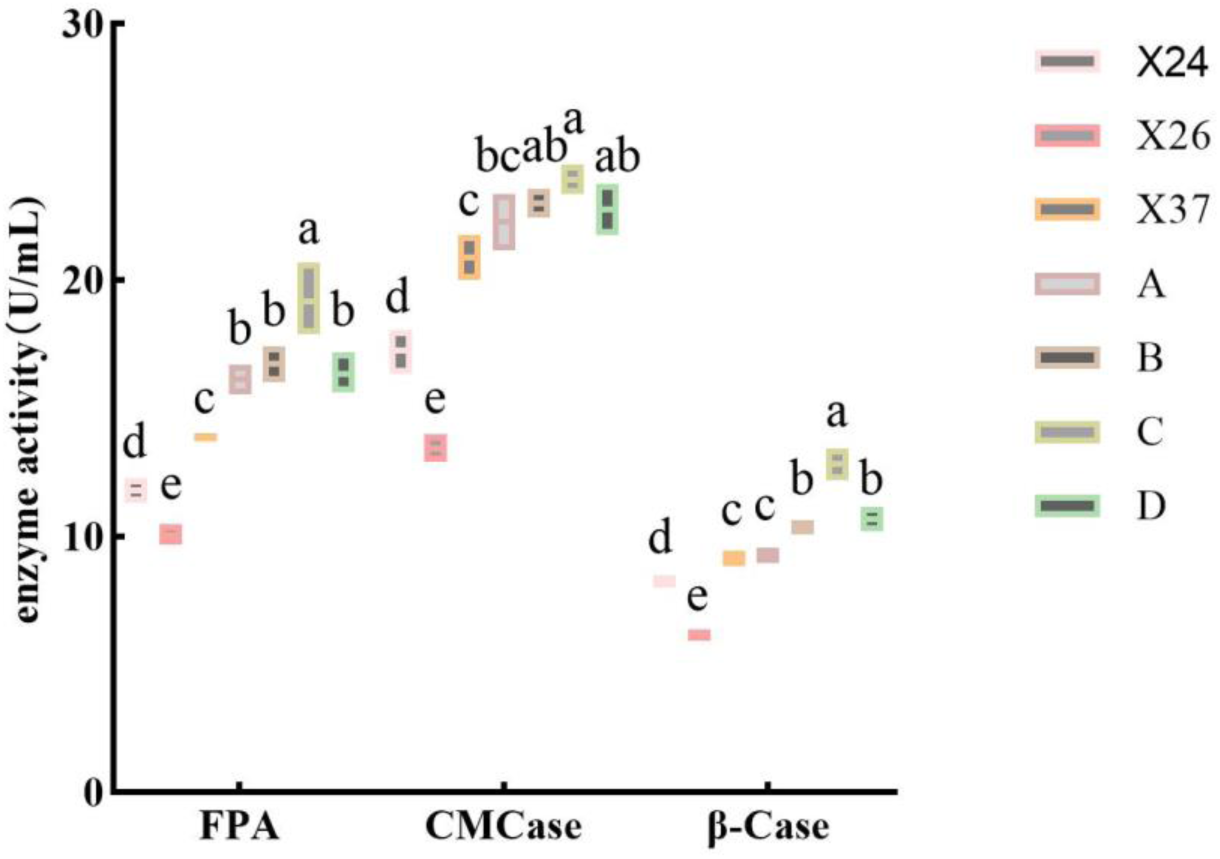
Enzyme activities of the three cellulose decomposing strain

### 3.5 Solid-liquid fermentation degradation rate

The CK group was the lank control group, and no bacterial solution was added. The weight loss rates of straw after solid fermentation and liquid fermentation were 15 % and 19 %, respectively. Compared with the CK group, the straw-degradation rate increased to different degrees in seven experimental groups: X24, X26, X37, A (X24+X26), B (X24+X37), C (X24+X26+X37) and D (X26+X37). It follows that the three strains of straw-degrading bacteria and their composite bacterial communities significantly promote the solid-liquid fermentation of corn straw. (Fig. 2). The results show that the solid-state fermentation degradation rate of the experimental group with the addition of strain *Pantoea rodasii* increased by 22 %, and the liquid-state fermentation degradation rate increased by 31 %, which was significantly higher than that by the other two single strain test groups, consistent with the results of enzyme activity determination. In all experimental groups, the community C increased the degradation rate of straw by 30 % in solid-state fermentation and 40 % in liquid-state fermentation, which was much higher than that by other groups, in accordance with the results of the enzyme activity measurements.

**Fig. 2.**
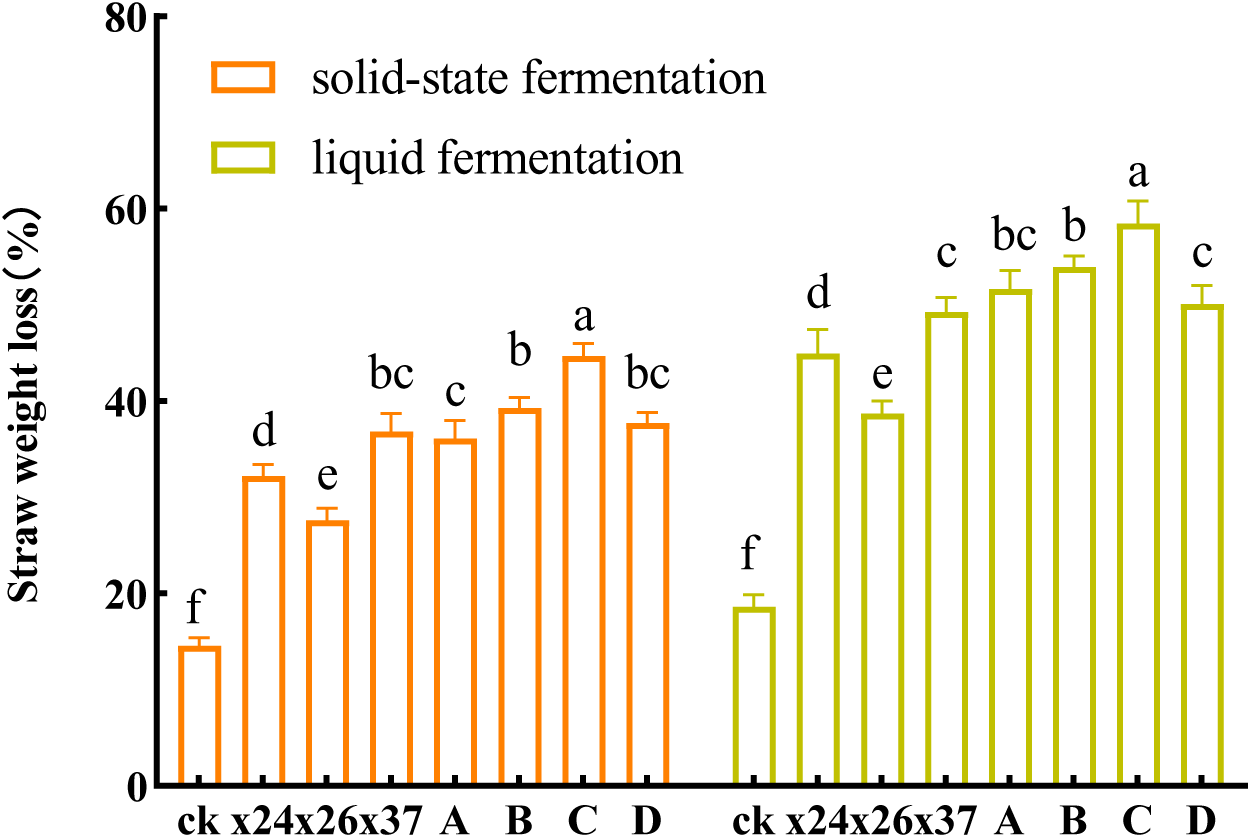
The results of straw liquid-solid fermentation

### 3.6 Optimization of enzyme production conditions of synthetic community C

The single-factor experiment was conducted on community C at different culture times, inoculation amounts, temperature, and pH to determine the optimal enzyme production conditions. The results show that the growth and CMCase activity of strain C were consistent with the change of different factors. The enzyme activity and growth of strain C increased first and then decreased with increasing culture time. When the culture time was 96 h, the CMCase activity reached a maximum value of 24 U / mL (Fig. 3a). When the inoculation amount was 4 %, the CMCase activity was the highest, 25 U / mL (Fig. 3b). At 12 ℃, the growth of community C was maximal, and the enzyme activity was the highest, 12 and 24 U / mL, respectively. The growth was faster at 16 ℃. It decreased rapidly at temperatures below 12 °C or higher than 16 °C, which shows that the temperature had a strong influence on strain C (Fig. 3c). The CMCase activity was the highest at pH 6.0, which was 24 U / mL (Fig. 3d). The results show that strain C had higher cellulase activity under neutral or weakly acidic conditions.

**Fig. 3.**
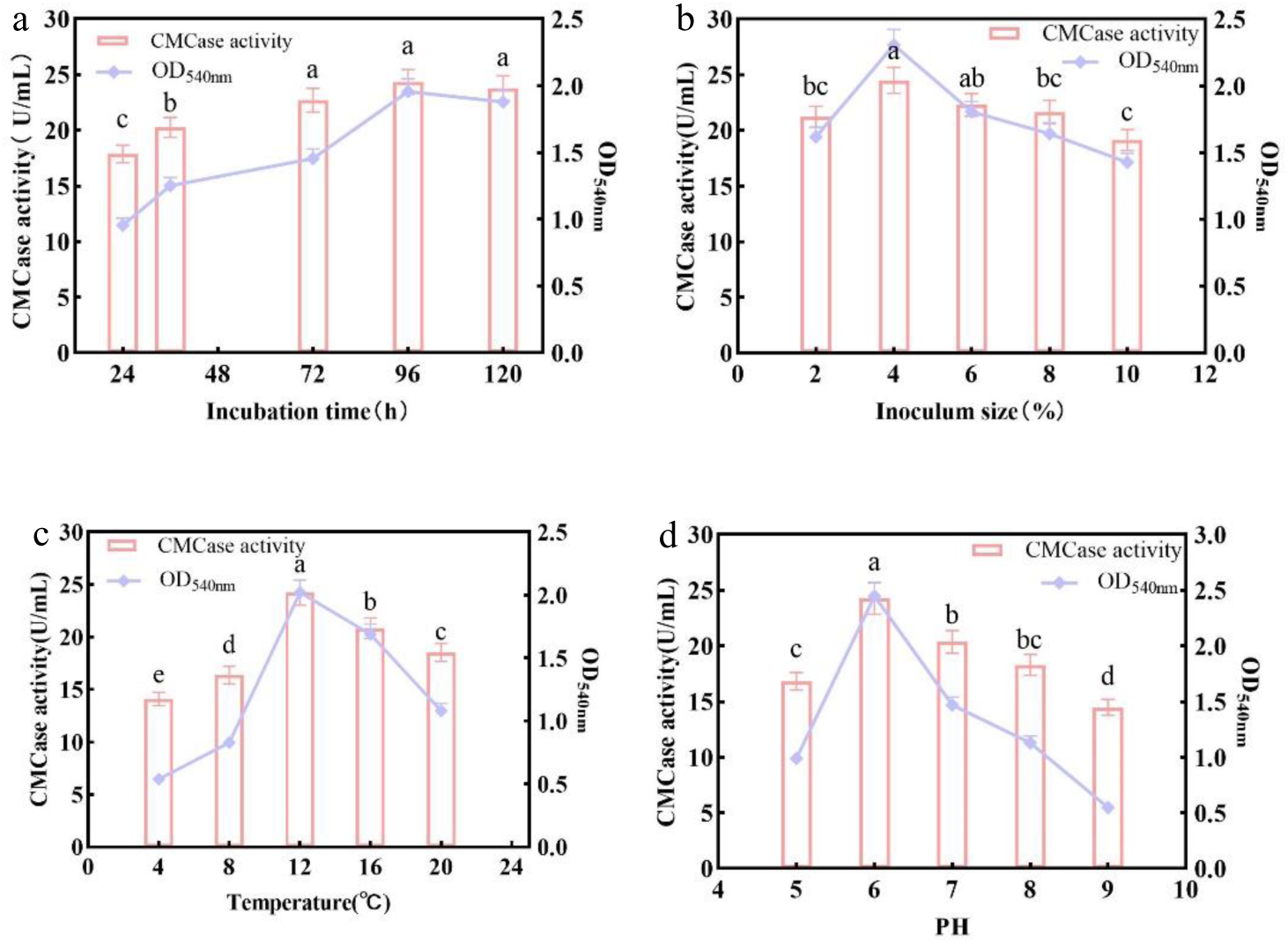
Effects of different factors on the growth and CMCase enzyme activity of complex strain C

## 4. Discussion

### 4.1 Screening of strains at low temperature

Isolation and screening of strains is a key step in the further study of molecular biological and biochemical characteristics such as strain identification and secretory enzyme determination (Zhao et al., 2018). Soil has always been a source of separation of various industrially-important microorganisms. Low-temperature limitation is a major factor inhibiting microbial action (Lü et al., 2012). In order to break the limitation, many have screened strains from the soil that still have the function of degrading cellulose at low temperature and constructed colonies, aiming to enhance the degradation rate of straw in winter. For example, the degradation rate of corn straw by degrading bacteria GF-20 can reach 32 % after 15 days of fermentation at 10 ℃ (Qinggeer et al., 2016). A consortium composed of *Bacillus subtilis, Pseudomonas* sp*., Citrobacter portucalensis*, *B. thringiensis, P. putida, Acinetobacter lwoffii* degrade 44 % of corn straw in 40 days at 15 ℃, which was 15 % more than the control (Han et al., 2023). No.1 composed of *Trichoderma* and various bacteria, and No.8 composed of *Penicillium* and various bacteria were cultured at 15℃ for 15 days, the decomposition rate of corn straw reached 30 % (Sa et al., 2013). The current low-temperature screening conditions were concentrated at 10 to 15℃. However, the winter soil temperature in Northeast China is lower than 10 ° C. Therefore, in this study, we conducted a low-temperature experiment using a straw returning surface soil sample that is rich in decomposing litter and bacterial diversity. The minimum induction temperature was 4 ℃. Finally, we selected three strains, *Stenotrophomonas maltophilia*, *Flavobacterium johnsoniae* and *Pantoea rodasii*, which have the ability to degrade straw efficiently at 4 to 20 ℃, ensuring that it has biological activity throughout the straw-returning process.

### 4.2 Analysis of straw degradation by dominant strains

In this study, we found that *Pantoea rodasii* had higher cellulolytic enzyme activity, and correspondingly higher cellulolytic capacity. In addition, *Pantoea rodasii* is a yellow rod-shaped Gram-negative bacterium in Enterobacteriaceae (Völksch et al., 2009). Initially, some researchers believed that *Pantoea rodasii* was a plant pathogen, and in some cases, it can secrete pathogenic factor extracellular polysaccharide (EPS) to promote plant pathogens to cause plant diseases (Roper, 2011). *Pantoea stewartii* can also colonize the xylem of plants, where it reproduces and produces EPS, blocking water flow and causing plants to wilt (Walterson & Stavrinides, 2015). Other researchers have identified epiphytic populations of *Pantoea rodasii* on many plants, including vegetables, fruits and grains, indicating that they have specific adaptations that promote phyllosphere colonization (Nadarasah & Stavrinides, 2014). This may also explain the higher lignocellulolytic capacity of *Pantoea rodasii*.

### 4.3 Analysis of straw degradation by bacterial communities

In the microbial community, the antagonism between microorganisms has a relatively stable negative impact on ecosystem functions (Steudel et al., 2016). In this study, antagonistic experiments of three strains were carried out, and the results show that there was no antagonistic effect between the three strains. The community C had the highest richness, including all three straw substrates with cellulose-degradation activity. There is a positive correlation between the richness of microbial communities and ecosystem functions (Wang et al., 2022), and the results of the present study also confirmed this. It is also consistent with the results of previous studies (Johnson et al., 2015; Loreau, 2010; Maron et al., 2018). Moreover, the positive effects of high abundance were mainly mediated by the number and types of community species (Wang et al., 2022). In this study, *Stenotrophomonas maltophilia* was first isolated from the soil of the Gangotri glacier. It can produce proteases at 5-25.8℃ and at a wide pH range (5-12), and the protease activity is not affected by repeated freezing and thawing (Kuddus & Ramteke, 2009). In community C, not only lignocellulose was decomposed, but also nitrogen-containing components in the straw were which improved the efficiency of the composite microbial system to degrade the straw. In addition, *Flavobacterium johnsoniae* also played an indispensable role in the community. A laminarinase was isolated and purified from the fermentation broth of *F. johnsoniae*, and the results show that it acted on the β-1,3-glycosidic bond of the soluble glucan substrate in an endoglucanase manner (Zhou et al., 2011). Some have pointed out that a way of microbial degradation of cellulose is from inside to outside, that is, straw crystalline cellulose is decomposed into amorphous cellulose soluble polysaccharide under the action of endoglucanase, and then decomposed into glucose by endoglucanase (Wang et al., 2016). In other words, *Flavobacterium johnsoniae* played a role in the decomposition of straw crystalline cellulose into amorphous cellulose. In addition, *F. johnsoniae* has the property of gliding along the surface of the substrate, and it has been hypothesized that there may be a relationship between the gliding property and the degradation ability of polysaccharides (McBride, 2004).

In conclusion, the results show that the degradation of cellulose by bacterial community C was significantly improved. Correspondingly, each strain in the community C played an important role in the degradation of straw. They may bear the function of decomposing crystalline cellulose in cellulose into non-qualitative cellulose, decomposing lignin and decomposing dry matter in straw, such as protein. Strains A, B and D showed that their overall effectiveness would be greatly reduced due to the random loss of one of the constituent strains.

## 4. Conclusions

We used a bottom-up strategy to construct a cold-tolerant corn straw-degrading community. First, we screened three strains, *Stenotrophomonas maltophilia*, *Flavobacterium johnsoniae*, and *Pantoea rodasii*, which produced cellulase with high activity at low temperatures, and they were not antagonistic to each other. Consequently, three highly-efficient strains were used to form a complex microbial system. After 45 days of fermentation at 12 ℃, the degradation rate of straw by strain C under solid-state fermentation was 45 %, 30 % higher than that of the control group. The degradation rate of liquid fermentation was 59 %, 40 % higher than the control group. The single-factor experiment was carried out on strain C, and the optimum conditions for enzyme production were determined as follows: culture time 96 h, inoculation amount 4 %, temperature 12 ℃, Ph 6.0. Under these conditions, the actual CMCase activity of strain C was 24 U / mL. In this study, a cold-tolerant corn straw-degrading community with high efficiency and stability was constructed, which is of great significance to the degradation of corn straw and the improvement of soil fertility in winter in Northeast China.

## Supplementary Information

Additional file 1: Table S1-S4.

## Acknowledgments

This work was supported by the National Key Research and Development Program of China (No. 2023YFD1502100), Basic Research Program of Liaoning Provincial Department of Education (No. JYTZD2023126).

